# Hindbrain V2a neurons impose rhythmic activity on motor neurons in an *in vitro* reticulospinal circuit

**DOI:** 10.1101/620674

**Authors:** Adele V Bubnys, Hagar Kandel, Lee-Ming Kow, Donald W Pfaff, Inna Tabansky

## Abstract

The reticulospinal system is an evolutionarily conserved pathway among vertebrates that relays locomotor control signals from the hindbrain to the spinal cord. Recent studies have identified specific hindbrain cell types that participate in this circuit, including Chx10^+^ neurons of the medullary reticular formation, which project to the spinal cord and are active during periods of locomotion. To create a system in which reticulospinal neurons communicate with spinal motor effectors, we have constructed an *in vitro* model using two purified excitatory neuronal subtypes: HB9^+^ spinal motor neurons and Chx10^+^ hindbrain neurons. Cultured separately, these neurons exhibit cell type-specific patterns of activity; the Chx10^+^ cultures developed regular, synchronized bursts of activity that recruited neurons across the entire culture, whereas motor neuron activity consisted of an irregular pattern. A combination of the two subtypes produced cultures in which Chx10^+^ neurons recruited the motor neurons into synchronized network bursts, which were dependent on AMPA receptors. In addition to demonstrating that the activity of *in vitro* networks can depend on the developmental identity of their constituent neurons, we provide a new model with genetically specified nerve cell types to study the activity of a reticulospinal circuit.

**Significance statement:** Models of the brain that use cultured neurons are usually comprised of a complex mixture of different kinds of cells, making it hard to determine how each cell type contributes to the overall pattern of activity. We made a simplified culture containing two cell types known to form a reticulospinal circuit *in vivo*. While in isolated cultures, each cell type had a distinct pattern of activity, in coculture the activity of one cell type came to dominate, indicating that the patterns observed in complex neuronal cultures arise in part from the distinctive properties of the constituent neurons.

## Introduction

To derive clinically relevant findings, most *in vitro* models of neurological disease seek to incorporate as realistic a mixture of cells from the modeled region as possible. These models of disease hold therapeutic promise, as some have already been used to identify small molecule candidates for drug development (Yang 2013, Wainger 2015, Ahfeldt 2017). Models of CNS regions have also been shown to develop complex patterns of activity similar to their *in vivo* counterparts (Trujillo 2018). Primary tissue taken from different areas of the brain and cultured on multi-electrode arrays (MEAs) develops regular bursts of spiking activity with tissue-specific differences in the shape of spike waveforms and the timing and structure of bursts (Dauth 2017, Soscia 2017, Sarkar 2018). When the different tissue types were co-cultured on a single array they developed correlated activity, suggesting that they could communicate with each other.

However, it is unclear whether these observations of bursts and changes in activity are an emergent property of the mixture of cell types in these cultures or governed by the presence of specific neuronal cell types (Maheswaranathan 2012, Chiappalone 2016, Lonardoni 2017). To appreciate how network activity emerges in such cultures and how it is determined by the properties of individual neuronal cell types, it is important to study each cell type in isolation prior to combining them into a more complex system.

To address this question, we created a simple system from two neuronal subtypes with a well-defined relationship in the intact nervous system, motor neurons and reticulospinal neurons. Motor neurons relay patterned input from spinal cord central pattern generator circuits to skeletal muscles to initiate behavior (Binder 1996). Reticulospinal neurons are components of a prominent behavioral circuit that relays rhythmic locomotor drive from the brainstem to the spinal cord (Peterson 1979, Garcia-Rill 1987a,b, Le Ray 2011, Kiehn 2016).

Of the subtypes of reticulospinal neurons, we specifically focused on V2a excitatory interneurons, identified by expression of the transcription factor Ceh-10 Homeodomain-Containing Homolog (Chx10, also known as Visual System Homeobox 2, or Vsx2), which are found in the spinal cord and medullary reticular formation. Chx10^+^ V2a neurons within the hindbrain reticular formation play a role in regulating hindlimb locomotion (Bretzner 2013, Bouvier 2015) and respiratory rhythm (Crone 2009, Crone 2012) and have functional connectivity to the mesencephalic locomotor region and pre-Bötzinger complex.

We hypothesized that the reticulospinal neurons cultured on MEAs would develop a different pattern of activity than motor neurons, which would be consistent with each subtype having a distinct behavioral role. Because motor neurons are controlled by reticulospinal neurons *in vivo*, we further hypothesized that the reticulospinal neurons’ activity would come to dominate in a combined coculture. This would support the idea that the electrical and biochemical properties of one neuronal subtype can drive the activity of the entire network *in vitro*.

Here, we report that Chx10^+^ hindbrain neurons develop synchronized network bursts that differ from the uncorrelated and irregular activity of motor neurons, and that in coculture motor neurons are recruited into Chx10^+^ neuron bursts. We then further identify some synaptic mechanisms that drive these circuit dynamics.

## Methods

### Cell culture

All cells were cultured at 37°C in 5% carbon dioxide and 95-100% humidity in Revco Ultima II CO_2_ incubators (Thermo Electron). Primary cortical glia were dissected and dissociated from Swiss Webster mice at P1-4 using the protocol described in Schlidge *et al*. (2013). Mouse pups were anesthetized with 5% isoflurane for five minutes, then decapitated. The forebrain was separated from the cerebellum and midbrain. The corpus callosum was severed, then the meningeal covering was peeled away. Forebrain tissue was dissociated in 10% trypsin (0.25% EDTA, Gibco, 25200-056) and passed through a 35μm filter (Corning, 352235). Cells were cultured on 100mm cell culture dishes treated with 0.1% gelatin (ATCC, PCS-999-027) at a density of ∼5×10^4^ cells/cm^2^ and grown until confluent, usually within 8 days. Glial culture media contained high glucose DMEM (Sigma-Aldrich, 51441C), 10% heat inactivated fetal bovine serum (ATCC, SCRR-30-2020), and 1% penicillin/streptomycin/antimycotic (Sigma-Aldrich, A5955). Once the glia reached confluence, they were dissociated with trypsin and cultured on sterile 5mm n.1 glass coverslips (Warner, 640700) treated with 1mg/ml Poly-D-Lysine (Millipore, A-003-E) and 1mg/ml laminin (Corning, 354232) in 24-well plates at a density of 5×10^5^ cells/well. Neurons were seeded on this feeder layer of glia once it reached confluence, after about 8 days.

ES-cell derived motor neurons were generated using the protocol described in Wichterle *et al*. (2002) from the HBG3 ES cell line, in which the enhanced green fluorescent protein (eGFP) is expressed under the control of the HB9 promoter (courtesy of Wichterle lab). ES cells were grown in ADFNK media that consisted of 1:1 DMEM/F12 (Millipore, DF-041-B): Neurobasal (Gibco, 21103049), 10% knock out serum replacement (Gibco, 10828010), 1% penicillin/streptomycin/antimycotic, and 1% GlutaMax supplement (Gibco, 35050061) for 2 days until they formed embryoid bodies. Media were supplemented on days 2 and 5 with 1μM retinoic acid (Sigma-Aldrich, R2625) and 1μM smoothened agonist (Calbiochem, 566661). On day 6, embryoid bodies were dissociated with papain according to manufacturer’s instructions (Worthington, LK003150).

Unsorted HB9^+^ motor neurons were plated on 5mm glass coverslips in a 24-well plate on top of a feeder layer of glia at a density of 1×10^6^ cells/well. HB9^+^ neurons that underwent FACS sorting were plated on Poly-D-lysine and laminin coated 5mm glass coverslips at a density of 5×10^5^ cells/well. For glial coculture, sorted HB9^+^ neurons were seeded on glass coverslips with a feeder layer of astroctyes at a density of 5×10^5^ cells/well. For multi-electrode recordings, standard 60-elecrode multi-electrode arrays (MultiChannel Systems, 890276) were sterilized and then coated with poly-D-lysine and laminin and seeded with 1×10^6^ sorted HB9^+^ neurons. For glial coculture on multi-electrode arrays, poly-D-lysine and laminin treated arrays were seeded with 5×10^5^ glial cells that were grown to confluence prior to seeding with 1×10^6^ sorted HB9^+^ neurons. Media consisted of the BrainPhys neuronal medium (StemCell, 5792) supplemented with 2% NeuroCult SM1 neuronal supplement (StemCell, 5711), 1% N2-supplement (Gibco, 17502048), 1% GlutaMax supplement, 1% pen/strep/antimycotic, 1μM Adenosine 3′,5′-cyclic monophosphate, N^6^,O2′-dibutyryl-sodium salt (dbCaMP, Calbiochem, 28745), 10ng/ml Brain derived neurotrophic factor (BDNF, MACS, 130-093-811), 10ng/ml Glial derived neurotrophic factor (GDNF, GoldBio, 1170-14-10), and 1μM ascorbic acid (Sigma-Aldrich, A4403). To produce HB9::GFP negative control for FACS sorting, ES-cell derived HB9^+^ motor neurons were generated in parallel from the E14 ES cell line (courtesy of Hatten lab).

Reticulospinal Ch×10^+^ neurons were dissected from E12.5 mouse embryonic hindbrains using the protocol described in Fantin *et al.* (2013) from mice in which the cyan fluorescent protein (CFP) is expressed under the control of the Chx10 promoter (Zhong *et al*, 2010). To produce the cells, a male mouse homozygous for Chx10::CFP (courtesy of Sharma lab) was mated with a Swiss Webster female mouse (Taconic). On day E12.5 of the pregnancy, the pregnant female was anesthetized in 5% isofluorane and oxygen and euthanized via cervical dislocation.

For the hindbrain dissection, each embryo was decapitated just rostral to the forelimb and the neural tube was isolated from the rest of the tissue. The developing rhombencephalon (hindbrain) segment corresponding to the position of the reticular formation in adults was excised and trimmed at the rostral and caudal ends. Dissections were performed in ice cold HBSS buffer (Gibco, 14175-095) supplemented with 1% pen/step/antimycotic, 20mM D-glucose (Sigma-Aldrich, G8769), and 1μM ascorbic acid. Hindbrains were dissociated with papain and sorted using flow cytometry to isolate the Ch×10^+^ subpopulation. To produce Ch×10::CFP negative control for FACS sorting, E12.5 hindbrains were derived from Swiss Webster mouse embryos. Sorted Ch×10^+^ hindbrain neurons were seeded on either 5mm glass coverslips in a 24-well plate or multi-electrode arrays, both prepared with a confluent layer of glia, at a density of 1×10^4^ neurons/well of coverslips or 4×10^4^ neurons/array. All Ch×10^+^ hindbrain neurons were cultured in Neurobasal medium supplemented with 2% SB-27 (Gibco, 17504044), 1% GlutaMax, 1% pen/strep/antimycotic, 1μM dbCaMP, 10ng/ml BDNF, 10ng/ml GDNF, and 1μM ascorbic acid.

For reticulospinal cocultures, sorted HB9::GFP^+^ motor neurons and Ch×10::CFP^+^ hindbrain neurons were seeded together on a confluent layer of glia on either 5mm coverslips or multi-electrode arrays. On coverslips in a 24-well plate, HB9^+^ neurons were seeded at a density of 2.5×10^5^ cells/well and Ch×10^+^ neurons were seeded at a density of 1×10^5^ cells/well. On multi-electrode arrays, HB9^+^ neurons were seeded at a density of 1×10^6^ cells/dish and Ch×10^+^ neurons were seeded at a density of 4×10^5^ cells/dish. Cocultures were grown in the same supplemented BrainPhys medium used for HB9^+^ cultures.

### Animals

Mice were group housed in a 12-hour light/dark schedule, with food and water provided *ad libitum*. For timed matings, two females were introduced into the home cage of a single male, where they remained for the duration of the mating. Females were checked for vaginal plugs every 24 hours and removed to separate cages after plug was detected, and singly housed for the duration of the timed pregnancy. All animal procedures and protocols were approved by the Rockefeller Institutional Animal Care and Use Committee.

### Flow Cytometry

All samples were sorted on the basis of fluorescent marker expression on the BD FACSAriaII benchtop flow cytometer with a 100μm nozzle and 20psi sheath pressure. Flow cytometry was performed at the Flow Cytometry Resource Center at Rockefeller University.

To isolate HB9::GFP^+^ motor neurons, embryoid bodies derived from HBG3 ES cells were dissociated on day 6 using papain and resuspended in FACS buffer for embryoid bodies that contains phenol-free HBSS supplemented with 2% heat-inhibited horse serum (Gibco, 26050088) and 5U/mL DNAse (Worthington, LK003172). For the GFP negative control, embryoid bodies were derived from E14 ES cells and prepared under parallel conditions. Between 10 and 20nM DAPI (Invitrogen, D1306) was added to each sample as a dead-cell exclusion dye. Each sample was excited by a violet 405nm laser and dead cells were excluded on the basis of emission in the DAPI wavelength 461nm using the 405D filter. Single cells were distinguished from doublets on the basis of forward and side scatter of the sample comparing the scatter area versus width. GFP fluorescence was detected using illumination from a 488nm blue laser equipped with a 535/30nm filter and the gate for GFP^+^ cell isolation was set based on a comparison of the GFP fluorescence of the HBG3-derived sample and the E14-derived sample. Typically, 50-60% of input cells from HBG3-derived embryoid bodies expressed GFP.

For Ch×10::CFP^+^ hindbrain neurons, hindbrains from Ch×10::CFP^+/−^ mice were dissociated at E12.5 using papain and resuspended in FACS buffer for hindbrains that contains high glucose phenol-free DMEM supplemented with 10% heat-inactivated fetal bovine serum, 1% pen/strep/antimycotic, and 5U/ml DNAse. For the CFP-negative control, hindbrains from Swiss Webster mice were prepared under parallel conditions. Approximately 20nM ToPro3 (Invitrogen, T3605) was added to each sample as a dead cell exclusion dye. Each sample was excited by a red 640nm laser, dead cells were excluded on the basis of emission in the ToPro3 wavelength using the 640C 670/30nm filter. As with the HB9::GFP^+^ motor neurons, single cells were distinguished from doublets on the basis of forward and side scatter area versus width. CFP fluorescence was detected using illumination from a 445nm blue violet laser equipped with a 490/30nm filter and the gate for CFP^+^ cell isolation was set based on a comparison of the CFP fluorescence of the Ch×10::CFP^+/−^-derived sample and the Swiss Webster-derived sample. Typically, 2.5-3% of input cells from Ch×10::CFP^+/−^ mouse hindbrains expressed CFP.

### Electrophysiology

Coverslips containing neurons cultured on a feeder layer of astrocytes as described above (see Cell Culture methods) for 5 to 10 days were perfused with 1× HEPES-ACSF in the recording chamber (HEPES-ACSF: 135mM NaCl, 10mM HEPES, 10mM glucose, 5mM KCl, 1mM CaCl_2_-2H_2_O, 1mM MgCl_2_) under constant flow (~5ml/minute). All cells were patched using pulled glass pipettes with an R_E_ of 5 to 12 MΩ filled with a standard internal pipette solution (K-gluconate: 14mM, HEPES-K: 10mM, NaHCO_3_: 60μM, Mg-ATP: 4mM, Na_2_-ATP: 2mM, Na-GTP: 30 μM, sucrose: 8mM, CaCl_2_: 1mM, EGTA: 5μM). Data were acquired on the MultiClamp 700B (Axon instruments) using ClampEx software. HB9^+^ spinal motor neurons were identified by GFP signal imaged using an Olympus BXS1W1 upright fluorescence microscope equipped with a FITC/EGFP filter (480/535nm ex/em, Chroma). Ch×10^+^ hindbrain neurons were identified by CFP signal from an ECFP filter (436/480nm ex/em, Chroma).

Once a giga-seal was achieved, the membrane voltage of the neuron was recorded for 1 minute at 1kHz sampling frequency without injecting additional current to measure the spontaneous activity of the neuron. For current-clamp experiments, current was injected to bring V_m_ to −70mV and current steps were applied in 10pA increments from −10 to 130pA for 1 second duration, returning to −70mV holding potential between steps. For voltage clamp experiments, the cell was held at −80pA for 100ms before stepping voltage injection from −100 to 150mV in 10mV increments for 100ms, returning to the −80pA holding potential between each step.

Data analysis and plotting of patch clamp data were performed using ClampFit and Matlab (see github.com/abubnys/patch_clamp_analysis for specific scripts used). To generate IV plots of voltage-gated sodium current from voltage-clamp data, the local minimum evoked current within 30ms of voltage step onset was subtracted from the mean current during the last 30ms of the voltage step and plotted against the magnitude of the injected voltage.

### Multi-electrode recordings

Multi-electrode arrays were cultured with HB9^+^ motor neurons or Ch×10^+^ hindbrain neurons as described above (see Cell Culture methods). For the duration of the lifetime of the culture (D3 to D30 days after plating for Ch×10^+^ and D7 to D30 days after plating for HB9^+^ neurons), spontaneous extracellular activity was recorded using the MEA2100-Lite system (MultiChannel Systems). The array was placed in the recording apparatus and allowed to equilibrate at room temperature for 30 minutes prior to recording for 4 minutes. Data acquisition was performed on MCRack with an input voltage range of −19.5 to +19.5mV and a sampling frequency of 20kHz. Raw electrode data for 60 electrodes were processed through a Bessel 4^th^ order high pass filter with a cutoff at 400Hz. The spike detection threshold was 5 standard deviations below the mean of the filtered recordings. Raw and filtered data, along with spike timestamps were converted to .txt files using MC_DataTool and the resulting files were analyzed in Matlab.

For wash-in experiments of the synaptic blockers 6-cyano-7-nitroquinoxaline-2,3-dione disodium salt (CNQX), D-(-)-2-amino-5-phosphonopentanoic acid (AP5), and bicuculline on the multi-electrode arrays, warmed 1× HEPES-ACSF was perfused through the MEA after the 30-minute equilibration period at ~5ml/minute for 10 minutes. The baseline activity of the MEA was recorded for 2 minutes under the previously mentioned parameters. Then, 100uL of a 10× solution of the drug was slowly perfused in at 50μL/minute for 2 minutes while recording. After 2 minutes the pump was stopped and the steady-state activity of the array in the presence of the drug was recorded for 4 minutes. The MEA was washed with 1× HEPES-ACSF at 5ml/minute for 10 minutes in between drug applications. Filtered electrode data were converted using MC_DataTool and analyzed in Matlab. The final concentrations of the drugs used were 20μM CNQX (Tocris, 479347-85-8), 50μM AP5 (Tocris, 79055-68-8), and 60μM bicuculline (Tocris, UN1544).

Data analysis for perfusion experiments was performed in Matlab (see github.com/abubnys/MEA_perfusion_package for specific scripts used). For initial spike data extraction, the high-pass filtered recordings from each electrode generated by MC_Rack (multichannel systems, Reutlingen, Germany) were converted into .txt format using MC_DataTool (multichannel systems). Spikes were detected in each channel using a manual threshold adjusted to pick up deviations that were approximately five standard deviations below the baseline of the recording and analysis of spike waveforms was used to determine whether one or more neurons was contributing to the observed signal. Spike sorting was performed on these data by plotting the aggregate collection of waveforms from recorded spikes. If this collection of waveforms fell within multiple visually distinguishable distributions, manual thresholds for each distribution were set by drawing a line through the waveforms visually classified as similar and then categorizing all recorded spikes according to whether they cross this threshold line or not. Then, spike rate was calculated for each waveform type by counting the number of spikes that fall within bins of 100ms width and multiplying by 10 to covert to units of Hz.

To facilitate comparison of different spike rates across all recordings, spike rates were smoothed using a spline function, binned according to the average spike rate in non-overlapping 10s intervals, then normalized to set the average spike rate from the first 10 bins (corresponding to the first 100s of recording) to 1.

To determine whether synaptic blocker wash-in had a dose-dependent effect on the activity of each culture type, normalized spike rate data from each electrode on the MEA that recorded spontaneous neuronal activity were pooled across all drug wash-in trials for a given culture type. The data corresponding to the period when the drug was washed in (2-4 minutes into the recording) were fit to a linear mixed effects model using the function fitlme() in Matlab with the normalized spike rate as the predictor variable and electrode as the random effect:

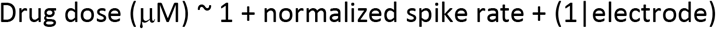

### Calcium imaging

HB9^+^, Ch×10^+^, or combined cocultures grown on 5mm coverslips with a feeder layer of glia were loaded with Rhod-3 AM dye according to the manufacturer’s instructions (Molecular Probes, R10145), then washed with 1× HEPES-ACSF. Calcium imaging was subsequently performed in 1× HEPES-ACSF.

For Ch×10^+^ cultures, calcium reporter dye fluorescence during spontaneous activity was imaged using an inverted spinning disc confocal microscope (Zeiss Axiovert 200) equipped with an EMCCD camera (Andor iXon). Solid state lasers were used for excitation at 443 and 561nm (Spectral Applied) paired with a polychroic filter with 440nm, 491nm, 561nm, and 640nm filters. Imaging acquisition was performed using MetaMorph software. Ch×10^+^ neurons were identified by CFP signal (440/480nm ex/em) and rhodamine3 signal was identified on the Texas Red channel (561/620-60nm ex/em). Calcium imaging data were acquired via time-lapse, with a 150ms interval and 100ms exposure time for 2 minutes.

For HB9^+^ cultures, HB9/Ch×10 cocultures, and astrocyte cultures, spontaneous calcium activity was imaged at room temperature and ambient CO_2_ using an Olympus BXS1W1 upright fluorescence microscope equipped with an Evolution QEi digital CCD camera (MediaCybernetics). A 120W mercury vapor short arc bulb was used as the fluorescence light source (X-Cite series 120Q). Imaging acquisition was done using NIS-Elements BR software. Hb9^+^ spinal motor neurons were identified by GFP signal using a FITC/EGFP filter (480/535nm ex/em, Chroma) and imaged with an exposure time of 100ms.

Ch×10^+^ hindbrain neurons were identified by CFP signal using an ECFP filter (436/480nm ex/em, Chroma) and imaged with an exposure time of 100ms. Rhodamine3 signal was imaged using a CY3/TRITC filter (545/605nm ex/em, Chroma) with an exposure time of 60ms per frame for 40 to 80 seconds.

For experiments involving application of the AMPA_R_ blocker CNQX, the spontaneous calcium activity of HB9/Ch×10 cocultures in HEPES-ACSF solution was imaged to determine a baseline level of activity. Then, 200μl of a 100× solution of CNQX was injected into the bath for a final drug concentration of 40μM. The culture was allowed to equilibrate for five minutes before imaging of spontaneous calcium activity in the presence of the drug. The drug was washed out by replacing 50% of media with fresh HEPES-ACSF in 5 repeated washes, then the culture was allowed to equilibrate for 5 minutes before measuring recovery of spontaneous activity.

Calcium imaging data for all experiments were analyzed in Matlab (see github.com/abubnys/calcium_imaging_ROI_analysis for specific scripts used). Due to overlap between CFP and GFP emissions spectra, CFP^+^ neurons appear on the GFP fluorescence channel and were distinguished from HB9::GFP^+^ neurons on the basis of their fluorescence on the CFP channel. ROIs were manually drawn around the cell bodies of identified CFP^+^ and GFP^+^ neurons and the mean Rhodamine3 fluorescence within the ROI was calculated at each frame of the recording in the Rhod3 channel.

Editing of rhodamine3 fluorescence time-course videos was performed in Fiji. For each video, brightness and contrast was adjusted uniformly across the image stack using the “auto” adjust function. Then, the minimum intensity for each pixel across the image stack (calculated using the “Z-project, minimum” function) was subtracted from each image in the stack to remove noise. Then, brightness and contrast were adjusted again across the image stack. Video playback is 20fps.

### Statistical Methods

All statistical analyses were performed in Matlab. Results are presented as mean ± SEM. For patch clamp experiments, age matched HB9::GFP^+^ neurons from cultures that were either immediately plated after dissociation from embryoid bodies or underwent flow cytometry prior to plating were subjected to the same battery of current clamp, voltage clamp, and spontaneous activity recordings. Student’s t-test was used for between-group comparisons of voltage-gated I_Na_ and spike threshold. For MEA recordings, cross-correlation was used to determine the degree of coordination across electrodes that detected spontaneous activity from each recording. For synaptic blocker experiments, all electrodes with spontaneous activity from MEA recordings were pooled according to culture type and normalized. To determine if there was a dose-dependent effect of synaptic blocker on spike rate, this data was fit to a linear mixed effects model. The statistical significance level for all of these analyses was set to p<0.05.

### Code accessibility

All custom written code used for this study is available on github. The code used to analyze and visualize patch clamp data is available at github.com/abubnys/patch_clamp_analysis. Code for quantifying calcium imaging data is available at github.com/abubnys/calcium_imaging_ROI_analysis. The code used for extracting and analyzing data from synaptic blocker perfusion experiments on MEAs is available at github.com/abubnys/MEA_perfusion_package.

## Results

### Developing reticulospinal cultures

Numerous studies of mixed populations of neurons from various brain regions including cortex, amygdala, and spinal cord have a strong tendency to develop network bursts when cultured on extracellular multi-electrode arrays. These bursts occur when many neurons across the cultured network fire at once at regular intervals (Wagenaar 2006, Black 2017, Dauth 2016, Van Pelt 2005). It is generally believed that the generation of such bursts requires a combination of excitatory and inhibitory interneurons that work in concert to balance network activity between a state of excitation and complete quiescence (Maheswaranathan 2012, Li 2009, Sternfeld 2017).

We sought to test this hypothesis by purifying neuronal subpopulations, which allowed us to culture identified neurons at defined stages of development. We focused specifically on reticulospinal cultures containing homogeneous populations of HB9^+^ spinal motor neurons and hindbrain Ch×10^+^ neurons. Hindbrain Ch×10^+^ neurons are known to play a role in regulating locomotor gait and breathing rhythm and they have descending projections to the spinal cord (Bretzner 2013, Bouvier 2015, Crone 2012). Spinal motor neurons provide direct limb muscle innervation. Thus, the *in vivo* function of both neuronal subtypes predisposes them to rhythmic bursts.

To isolate pure populations of HB9^+^ spinal motor neurons and hindbrain Ch×10^+^ neurons, we employed fluorescence activated cell sorting (FACS). We cultured these cell types as single populations and also as a mixed reticulospinal culture. We differentiated HB9^+^ spinal motor neurons from HBG3 embryonic stem cells using Wichterle et al’s protocol to induce the spinal motor neuron identity (Wichterle 2002). Embryoid bodies were dissociated 6 days after formation and sorted on the basis of HB9::GFP expression. E14 stem cells lacking GFP were used as a negative control for FACS (Figure 1b,c). Approximately 50-60% of unsorted cells in the embryoid body derived from HBG3 ES cells expressed GFP. FACS sorting for GFP expression enriched this population to >96% purity. HB9::GFP^+^ motor neurons were subsequently cultured on a layer of cortical astrocytes to improve axonal outgrowth and network development.

**Figure 1:**
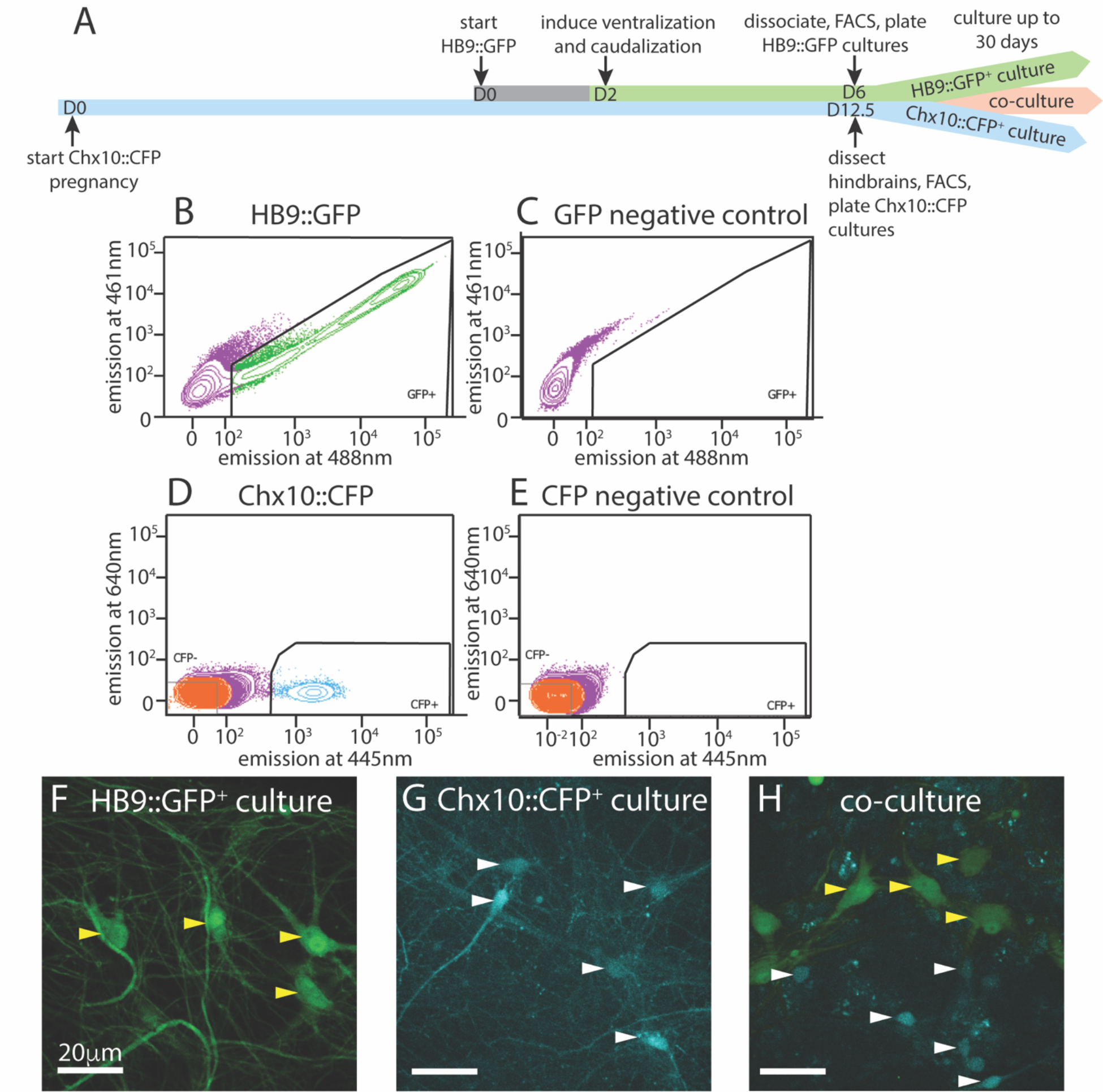
Isolation and culture of HB9^+^ motor neurons and Ch×10^+^ hindbrain neurons. ***A***, Timeline schematic of isolating and setting up HB9::GFP^+^, Ch×10::CFP^+^, and combined co-cultures. **B-E** Sample FACS plots and thresholds for isolation of Hb9::GFP^+^ and Ch×10::CFP^+^ neurons. ***B**,* GFP^+^ neurons from HB9::GFP stem-cell derived embryoid bodies after 6 days in culture (DIC) ***C**,* embryoid bodies derived from non-transgenic ES cells (negative control) ***D**,* CFP^+^ neurons from E12.5 hindbrains of Ch×10::CFP mice and ***E**,* Swiss Webster mice (negative control). **F-H** Fluorescent photomicrographs of neurons cultured after sorting. Yellow arrowheads indicate HB9::GFP^+^ neurons and white arrowheads indicate Ch×10::CFP^+^ neurons. ***F***, Sorted HB9::GFP^+^ neurons, 16 DIC. ***G***, sorted Ch×10::CFP^+^ hindbrain neurons, 10 DIC (scale bar 20μm) ***H***, combined culture of both subtypes, 16 DIC (scale bar 20μm).

To test whether sorting affected the electrophysiological activity of HB9^+^ neurons, we performed whole cell patch clamp on HB9::GFP^+^ neurons from sorted and unsorted cultures grown in parallel under identical conditions. After 7 days in culture, HB9^+^ neurons in both treatments responded to brief current pulses with spike trains, having a spike threshold around 20pA (Figure 2a,b). They developed voltage gated sodium current (I_Na_) with maximum current evoked at −2±12 mV (Figure 2c-e) that was not significantly different between sorted and unsorted populations (Student’s 2-tailed T-test p = 0.879). After 13 days in culture, both sorted and unsorted HB9^+^ motor neurons also developed spontaneous spike trains (Figure 2f).

**Figure 2:**
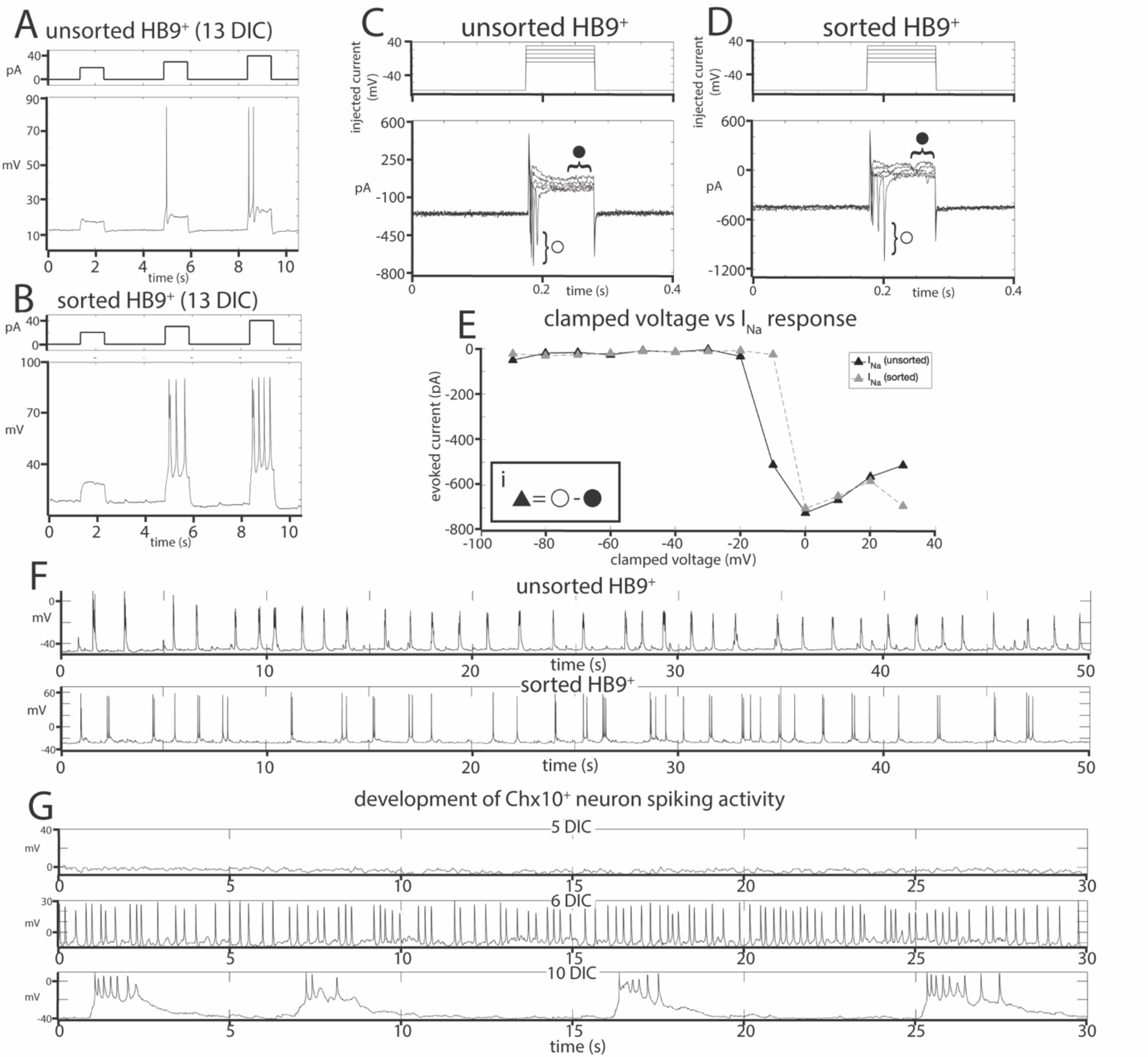
Effects of FACS sorting on neuron electrophysiology. **A-B** Comparison of the response of ***A***, unsorted and ***B***, sorted HB9::GFP^+^ neurons (bottom panels) to injections of 20, 30, and 40pA current (top panels). **C-D** Response of ***C***, unsorted and ***D***, sorted HB9::GFP^+^ neurons (bottom panels) to voltage step injection of −90 to 30mV (top panels, result for −10 to 30mV injections shown) (7 DIC). Sodium current (I_Na_) was calculated at each injected voltage step by subtracting the steady state current response (•) from the initial current minimum (o) (formula shown in insert *Ei*). ***E***, I-V plot of Na^+^ currents for sorted and unsorted HB9::GFP^+^ neurons calculated from the voltage clamp experiment results shown in *C* and *D*. ***F**,* Spontaneous activity of unsorted (top panel) and sorted (bottom panel) HB9::GFP^+^ cells at 13 DIC***. G***, Spontaneous activity of sorted Ch×10::CFP^+^ neurons at 5 DIC (top), 6 DIC (middle) and 10 DIC (bottom).

We then isolated and cultured primary hindbrain neurons expressing the transcription factor Ch×10, also using the FACS approach. We first assessed the Ch×10^+^ neurons’ behavior *in vitro* as a homogeneous population, and then in combination with HB9^+^neurons to determine if they could form a reticulospinal circuit *in vitro*. For these experiments, we dissected neurons from embryonic Ch×10::CFP^+/−^ mice at E12.5, prepared a single cell suspension and used FACS to isolate the CFP^+^ population. As a negative control for CFP expression, we used hindbrains taken from wildtype (WT) Swiss Webster E12.5 mouse embryos that do not express CFP (Figure 1d,e). The hindbrains contained 2-3% Ch×10::CFP^+^ neurons, and sorting enriched this population to >95% purity. These CFP^+^ neurons were then cultured on a layer of cortical astrocytes, which is known to improve the development and long-term viability of neuronal cultures (Wang 1994, Maher 1999, Boehler 2004).

It is possible that, when removed from the intact reticular formation with its descending inputs and diversity of other cell types, Ch×10^+^ hindbrain neurons would not develop any intrinsic activity that could pattern a reticulospinal circuit. To assess the electrophysiological development of sorted Ch×10^+^ neurons, we used whole-cell patch clamp to record the spontaneous activity of single cells in cultures at different ages ranging from 1 to 30 days in culture. For Ch×10^+^ hindbrain neurons, the measured membrane capacitance was 22.75±2pF, membrane resistance was 787.27±105MΩ, access resistance was 29.01±3MΩ, and membrane voltage was −22.6±4mV. We found that Ch×10^+^ hindbrain neurons developed spontaneous electrophysiological activity after 5 days in culture. This activity started off as random trains of spikes, but gradually became organized into robust, regular bursts by 10 days in culture and this pattern of activity continued throughout the remaining lifetime of the cultures (Figure 2g).

### Motor and Ch×10 neuron cultures develop distinct patterns of network activity

Having established that HB9^+^ motor neurons and Ch×10^+^ hindbrain neurons develop spontaneous electrophysiological activity at the single cell level, we sought to determine whether cultures of either cell type, which are composed almost exclusively of excitatory neurons and astrocytes, could generate spontaneous patterns of network activity, whether these patterns would organize into network bursts, and whether there were any cell-type specific differences in such activity.

To record the activity of multiple neurons at different time points, we cultured sorted HB9^+^ motor neurons on multielectrode arrays (MEAs) containing a grid of 64 extracellular recording electrodes. We recorded their spontaneous activity daily over 30 days, starting from the day after plating.

We found that on their own, without astrocytes, sorted HB9^+^ motor neurons did not develop any spontaneous activity on the MEA (n = 6). However, when these neurons were cultured on a confluent layer of astrocytes, they gradually developed robust network activity that remained stable over a month of recording (n = 14). We note that astrocytes cultured on their own did not develop spontaneous activity when recorded on MEAs (n = 3), although we did observe spontaneous calcium flux in astrocyte cultures visualized with the calcium-sensitive dye Rhodamine3 (**Video 3-1**). The activity of HB9^+^ motor neuron/astrocyte cultures was not well coordinated, even among neighboring recording electrodes (Figure 3a-c). To assess whether the overall activity of the culture had a hidden underlying temporal structure, we calculated the mean spike rate across all active channels of the HB9^+^ motor neuron cultures and found that it remained constant throughout the recording session (Figure 3d).

**Figure 3:**
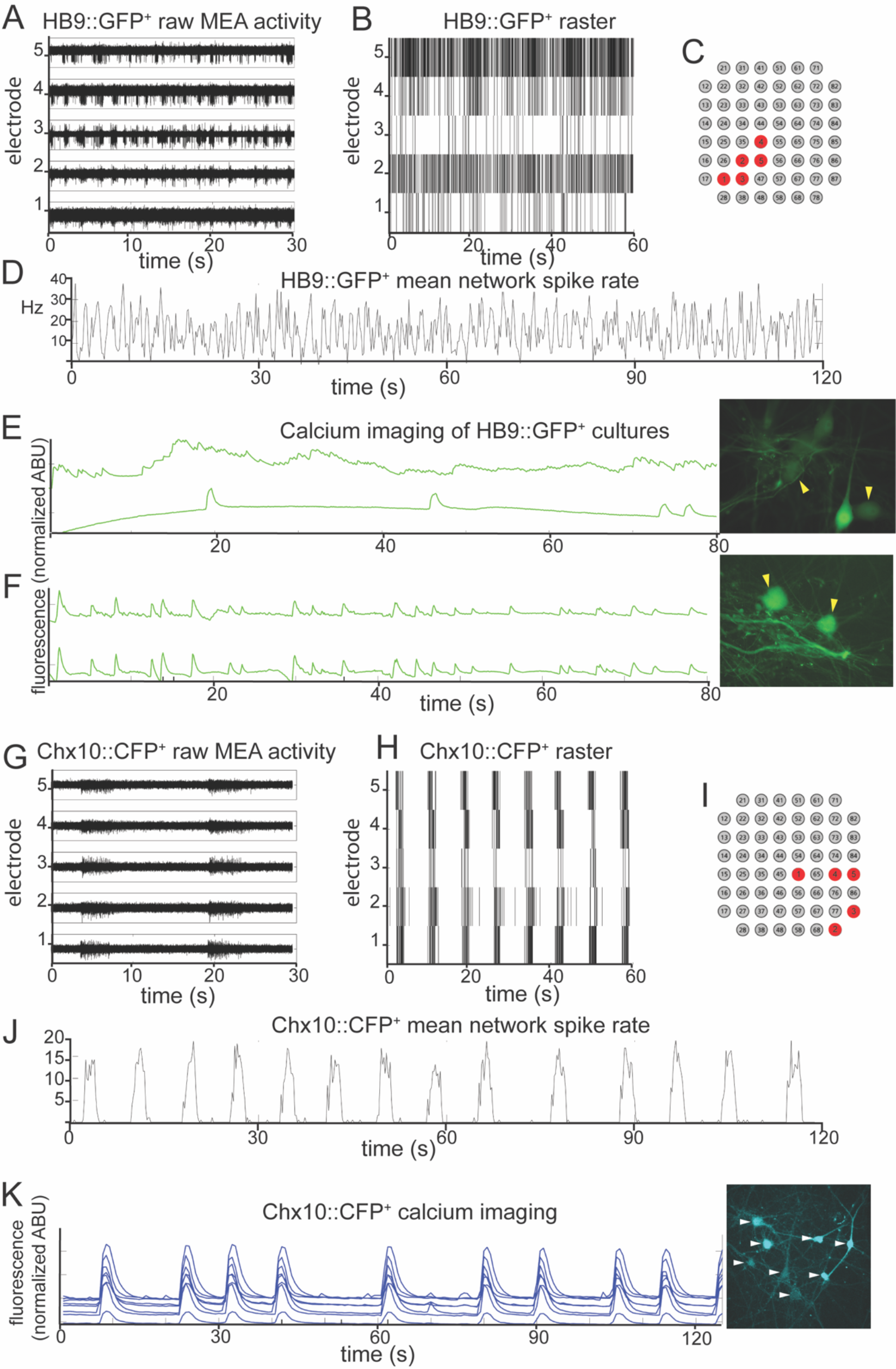
HB9^+^ motor neurons and Ch×10^+^ hindbrain neurons develop different patterns of activity *in vitro*. **A-C** Example of a multielectrode array (MEA) recording of sorted HB9::GFP^+^ neurons (18 DIC), both as ***A***, high pass filtered MEA data and ***B***, raster plot. Locations of electrodes indicated in red in ***C***. ***D***, Mean spike rate of entire HB9::GFP^+^ neuron culture from A,B. **E-F** Quantification of calcium-sensitive Rhodamine3 dye fluorescence in the cell bodies of HB9::GFP^+^ neurons ***E***, 19 DIC, see video 3-2 for rhodamine fluorescence time course, ***F***, 32 DIC, see video 3-3. Corresponding right panels are photomicrographs of neurons quantified for calcium activity, indicated by yellow arrowheads. **G-I** Example of a multielectrode array (MEA) recording of sorted Ch×10::CFP^+^ neurons (5 DIC) as ***G***, high-pass filtered MEA data and ***H***, raster plot. locations of electrodes on array in red in ***I***. ***J***, Mean spike rate of entire Ch×10::CFP^+^ neuron culture from B,E. ***K***, Calcium imaging of Ch×10::CFP^+^ neurons (10 DIC), also shown in video 3-4. Right panel is a photomicrograph of identified neurons quantified for calcium activity, indicated by white arrowheads.

When we used the calcium-sensitive dye Rhodamine3 to assess HB9^+^ motor neuron activity with single-cell resolution, we observed randomly distributed calcium spikes that were asynchronous between neighboring neurons (Figure 3e, **Video 3-2**), though more mature cultures did develop some synchrony (Figure 3f, **Video 3-3**). The mean correlation coefficient between the spike rates of multiple neurons within the same HB9^+^ neuron culture was 0.15±0.17 (p = 0.15).

When we cultured Ch×10^+^ hindbrain neurons on MEAs with a confluent layer of astrocytes, we observed the emergence of spontaneous activity with these neurons as well. Unlike HB9^+^ neurons, Ch×10^+^ neurons developed robust and coordinated network bursts (Figure 3g-j). Practically no spikes occurred outside of these sharply delineated bursting periods. The time between bursts (inter-burst interval) varied between 2 and 10 seconds throughout the lifetime of the cultures, with no apparent long-term trend. We observed the same sort of robust network bursts in Ch×10^+^ hindbrain neuron cultures with calcium imaging (Figure 3k, **Video 3-4**).

### Ch×10^+^ neurons impose their activity patterns on HB9^+^ neurons in coculture

Despite their common glutamatergic identity, we observed that HB9^+^ and Ch×10^+^ hindbrain neurons develop distinct patterns of spontaneous network activity. If these two cell types fail to form functional connections to one another *in vitro*, these patterns of activity should remain unchanged in coculture, but if a unidirectional functional connection forms between Ch×10^+^ and HB9^+^ neurons, we might expect to see one activity pattern dominate in coculture. To test these possibilities, we cultured the two cell types together as a mixed population on MEAs and recorded their spontaneous activity daily over 30 days. Such cocultures develop spontaneous bursts of comparable time scale and duration to pure Ch×10^+^ cultures, though some neurons continue to have spiking activity that resembles HB9^+^ motor neurons in between network bursts (Figure 4a-c). When the overall network activity was measured by averaging spike rates across all active electrodes, the Ch×10-like network bursts predominated (Figure 4d).

**Figure 4:**
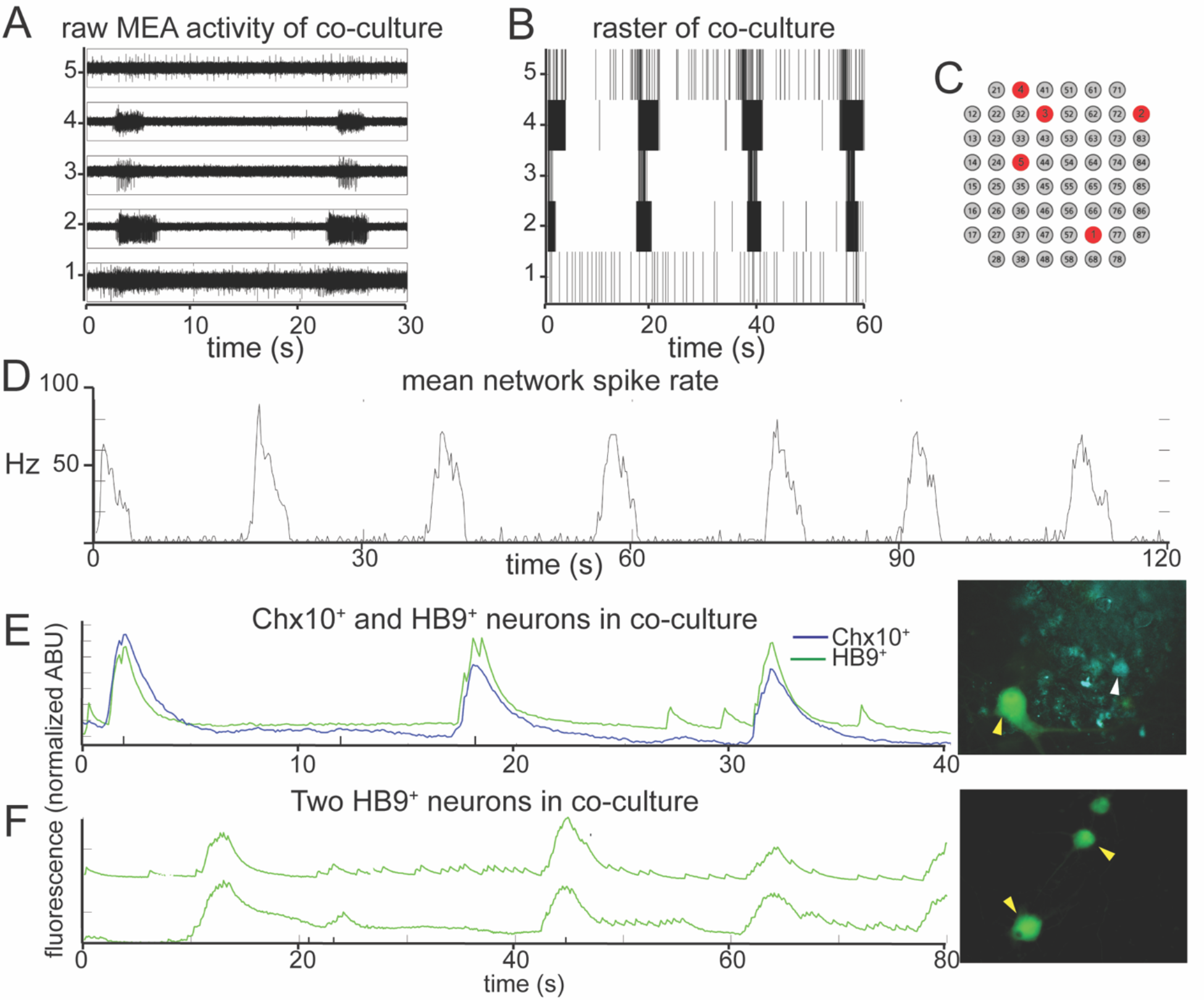
In reticulospinal co-culture, Ch×10^+^ hindbrain neurons drive patterned HB9^+^ neuron activity. **A-B** Example of an MEA recording of HB9::GFP^+^/Ch×10::CFP^+^ neuron coculture (8 DIC), both as ***A***, high pass filtered MEA data and ***B***, raster plot. Locations of electrodes indicated in red in ***C***. ***D***, Mean spike rate of entire co-culture from A, B over the course of 120 seconds. **E-F** Calcium imaging of neurons in co-culture. ***E***, Normalized calcium-sensitive fluorescence intensity over time in co-cultured HB9::GFP^+^ and Ch×10::CFP^+^ neurons participating in coordinated bursts, shown also in video 4-1. ***F***, Normalized calcium-sensitive fluorescence intensity of two HB9::GFP^+^ neurons from co-culture (Ch×10::CFP^+^ neurons not pictured) participating in network bursts, shown also in video 4-2. Corresponding right panels indicate identified neurons quantified for calcium activity, white arrowheads for Ch×10::CFP^+^ and yellow arrowheads for HB9::GFP^+^.

It is possible that the bursts we observed in the reticulospinal culture were generated only by the Ch×10^+^ neurons in the dish and that the HB9^+^ motor neurons were quiescent and did not contribute to network activity. In order to determine which cell type participates in the cultures’ network bursts, we used calcium imaging to obtain single cell resolution recordings of the coculture. We found that neighboring HB9^+^ and Ch×10^+^ neurons both participate in network burst events (Figure 4e, **Video 4-1**). Some HB9^+^ motor neurons in coculture also have brief, non-coordinated calcium spiking events that occur between the larger bursts (Figure 4f, **Video 4-2**).

The percentages of Ch×10^+^ and HB9^+^ neurons from calcium-imaging experiments that were spiking, bursting, both spiking and bursting, or inactive in each of the culture conditions are summarized in Table 1.

**Table 1:** Overview of activity patterns of Ch×10^+^ and HB9^+^ neurons from calcium imaging experiments.

| Cell type | Number total cells recorded | Spiking: number (percentage) | Bursting: number (percentage) | Spiking and Bursting: number (percentage) | No activity: number (percentage) |
| --- | --- | --- | --- | --- | --- |
| **Chx10<sup>+</sup> (co-culture)** | 125 | 16 (13%) | 64 (51%) | 17 (14%) | 28 (22%) |
| **Chx10<sup>+</sup> (alone)** | 483 | 0 | 312 (65%) | 7 (1%) | 164 (34%) |
| **HB9<sup>+</sup> (co-culture)** | 67 | 13 (19.5%) | 13 (19.5%) | 29 (43%) | 12 (18%) |
| **HB9<sup>+</sup> (alone)** | 176 | 81 (46%) | 0 | 2 (1%) | 93 (53%) |

### HB9^+^ and Ch×10 network activity is an AMPA receptor-dependent process

The spontaneous coordinated activity we observed in Ch×10^+^ and HB9^+^ neuron cultures could be the product of intrinsic pacemaker properties of these neurons or an emergent property of the network that is dependent on synaptic transmission. To distinguish between these alternatives, we applied a panel of synaptic blockers targeting α-amino-3-hydroxy-5-methyl-4-isoxazolepropionic acid (AMPA) receptors, *N*-methyl D-aspartate (NMDA) receptors, and γ-aminobutyric acid, type A (GABA_A_) receptors, while recording from the cultures on MEAs to observe changes in spontaneous activity. The blockers used included the AMPA receptor antagonist 6-cyano-7-nitroquinoxaline-2,3-dione disodium salt (CNQX), the NMDA receptor antagonist D-(-)-2-amino-5-phosphonopentanoic acid (AP5), and the GABA_A_ receptor antagonist bicuculline. Washing in the AMPA_R_ antagonist CNQX on cultures of spiking HB9^+^ neurons caused a gradual decrease in activity to about 40% of initial levels (Figure 5a, e). There was a significant relationship between drug dose and spike rates (linear mixed effects model: β = −0.04, p = 2.65e-63). Similarly, CNQX application resulted in a significant decrease in the activity of Ch×10^+^ neurons to about 40% of the initial rate (Figure 5b, f) (β = −0.021, p = 8.61e-15). The application of CNQX to cocultures caused the majority of cells to abruptly stop bursting (Figure 5c). Other neurons gradually became decoupled from the network bursts and fired tonically for a brief period before also being silenced during CNQX application (figure 5d). The average response of cocultured neurons to CNQX application reflects this transient increase in activity followed by eventual inhibition (Figure 5g) (β = −0.012, p = 0.0015).

**Figure 5:**
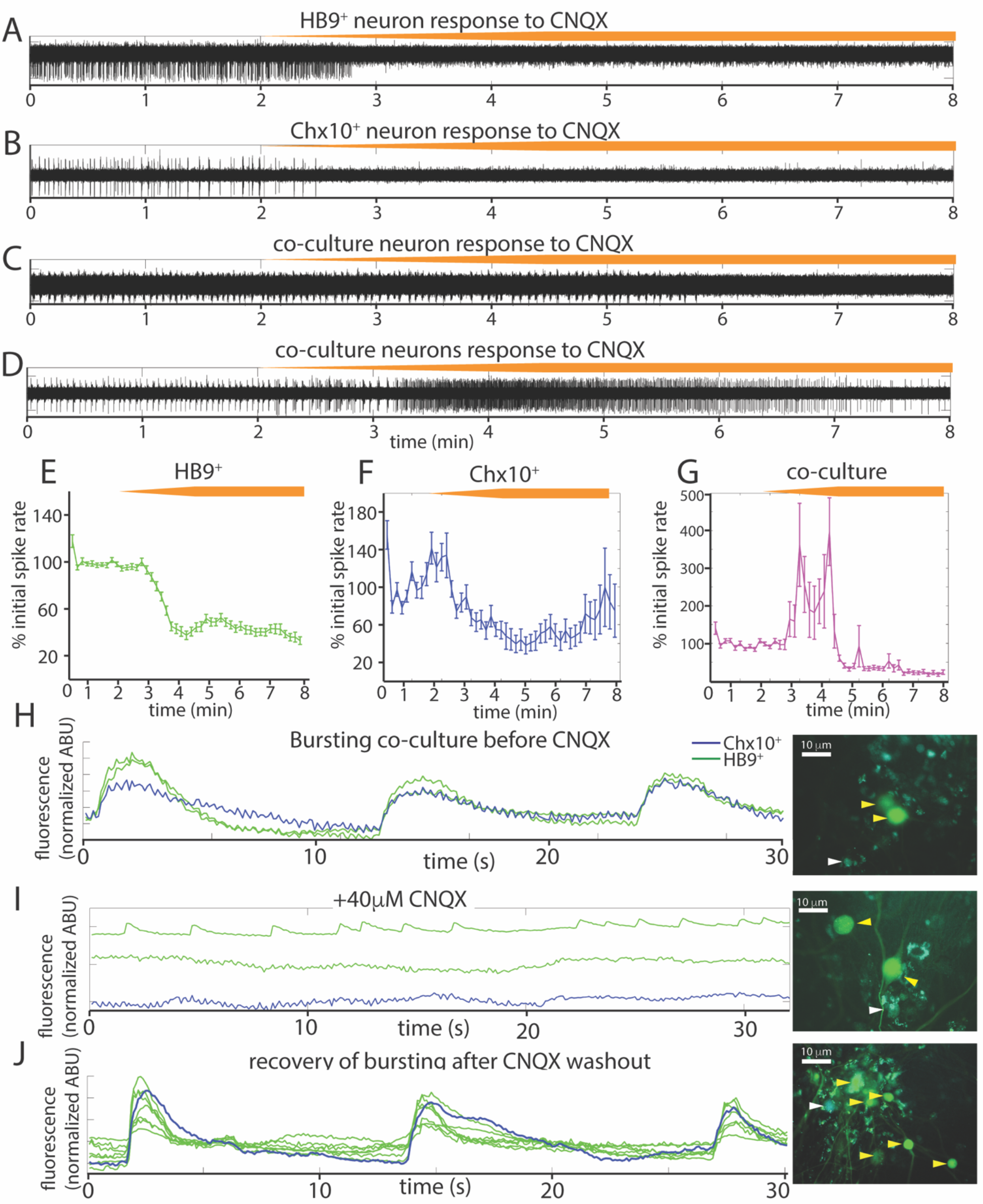
Spontaneous activity in reticulospinal cultures is an AMPA_R_-dependent process. **A-D**, Examples of high pass filtered MEA recordings of spiking neurons during wash-in of a 200μM solution of the AMPA_R_ blocker CNQX at 50μL/min (final CNQX concentration 20μM), orange bars show time course of blocker wash-in. ***A***, Neuron from HB9::GFP^+^ culture, ***B*** neuron from Ch×10::CFP^+^ culture, ***C-D*** examples of two different kinds of responses to CNQX of neurons from HB9::GFP^+^/Ch×10::CFP^+^ co-culture. **E-G**, Normalized mean responses of all neurons recorded from electrodes with activity to CNQX wash-in, ***E*** HB9::GFP^+^ cultures (n = 3), ***F*** Ch×10::CFP^+^ cultures (n = 3), ***G*** HB9::GFP^+^/Ch×10::CFP^+^ co-cultures (n = 4). **H-J**, Calcium imaging of co-culture ***H***, bursting prior to CNQX application (shown also in video 5-1), ***I***, inhibition of bursting, but not Hb9::GFP^+^ spiking, by application of 40 μM CNQX (shown also in video 5-2), and ***J***, bursting recovers after washout of CNQX (shown also in video 5-3). Corresponding right panels are photomicrographs of neurons quantified for calcium activity, indicated by white arrowheads for Ch×10::CFP^+^ and yellow arrowheads for HB9::GFP^+^.

We repeated the CNQX drug application on cocultures and used calcium imaging with Rhodamine3 to visualize the activity of the culture prior to and after application of 40μM CNQX. Despite a loss of network bursting activity, we observed that some HB9^+^ neurons in the coculture continued to have spontaneous spiking activity in the presence of a blocking concentration of CNQX (Figure 5h-j).

We also tested the effects of the NMDA receptor antagonist AP5 on all three cultures (Figure 6a-f) and found that there was no significant relationship between blocker dose and spike rates during AP5 wash-in (linear mixed effects model for: HB9^+^ neurons, β = 0.0005 p = 0.23, Ch×10^+^ neurons, β = 0.004 p = 0.25, coculture, β = 0.006 p = 0.24). The GABA_A_ receptor blocker bicuculline also had no detectable effect on Ch×10^+^ hindbrain neurons, HB9^+^ motor neurons, or cocultures (Figure 6g-l) (linear mixed effects model for: HB9^+^ neurons, β = 0.0003 p = 0.54, Ch×10^+^ neurons, β = 0.0026 p = 0.34, coculture,β = 0.0057 p = 0.13).

**Figure 6:**
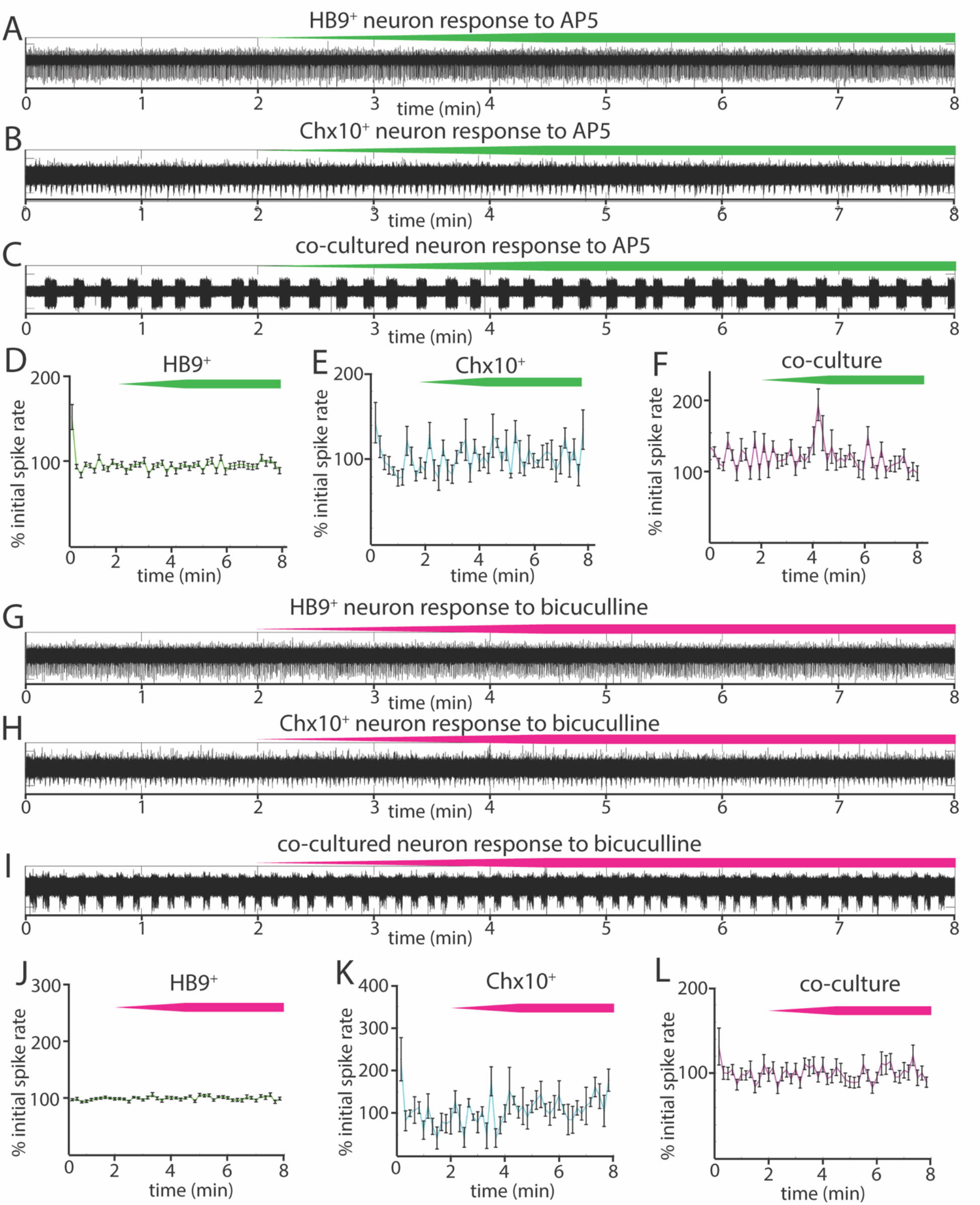
Responses of reticulospinal cultures to NMDA_R_ and GABA_A_R blockers. **A-C** Examples of high pass filtered MEA recordings of spiking neurons during wash-in of a 500μM solution of NMDA_R_ blocker AP5 at 50μL/min (final AP5 concentration 50μM), green bars show approximate time course of AP5 wash-in. ***A***, Neuron from HB9::GFP^+^ culture, ***B*** neuron from Ch×10::CFP^+^ culture, ***C*** neuron from HB9::GFP^+^/Ch×10::CFP^+^ coculture. **D-F**, Normalized mean responses of all recorded neurons to AP5 wash-in, ***D*** HB9::GFP^+^ cultures (n = 3), ***E*** Ch×10::CFP^+^ cultures (n = 3), ***F*** HB9::GFP^+^/Ch×10::CFP^+^ co-cultures (n = 4). **G-I** Examples of high pass filtered MEA recordings of spiking neurons during wash-in of a 600μM solution of GABA_A_R blocker bicuculline at 50μL/min (final bicuculline concentration 60μM), magenta bars show time course of bicuculline wash-in. ***G***, Neuron from HB9::GFP^+^ culture, ***H*** neuron from Ch×10::CFP^+^ culture, ***I*** neuron from HB9::GFP^+^/Ch×10::CFP^+^ coculture. **J-L** Normalized mean responses of all recorded neurons to bicuculline wash-in, ***J*** HB9::GFP^+^ cultures (n = 3), ***K*** Ch×10::CFP^+^ cultures (n = 3), ***L*** HB9::GFP^+^/Ch×10::CFP^+^ cocultures (n = 4).

## Discussion

In this study, we used flow cytometry to isolate HB9^+^ motor neurons and Ch×10^+^ hindbrain neurons and cultured these cell types separately and together to form a reticulospinal circuit. We found that the sorting process did not significantly impact the development of HB9^+^ and Ch×10^+^ neuron electrophysiology. When isolated, these two cell types developed distinct patterns of network activity. HB9^+^ neurons tended towards uncoordinated spike trains, while Ch×10^+^ hindbrain neurons were characterized by regular, network-wide bursts of activity. Cocultures of these two cell types developed the network bursts characteristic of Ch×10^+^ neurons that recruited neighboring HB9^+^ neurons. We further note that the activity of all these cultures was insensitive to NMDA and GABA_A_ receptor blockers but could be inhibited by the AMPA receptor blocker CNQX.

### Effect of cell sorting on electrophysiology of isolated cells

FACS-sorted stem-cell derived HB9^+^ motor neurons develop complex morphology and electrical excitability *in vitro* (Uzel 2016, Yang 2013, Haidet-Phillips 2011, Wichterle 2002), but previous studies had not established whether the nature of their electrical responses had been altered. Our results show that sorted Hb9^+^ motor neurons develop spontaneous spiking activity, fast inactivating sodium currents, and repetitive trains of action potentials in response to current injection. These results are consistent with the reported electrophysiology of unsorted stem cell-derived motor neurons (Miles 2004). Thus, our single-cell electrophysiology indicates that the presence of other neuronal subtypes and progenitors does not alter the electrical properties of HB9^+^ motor neurons, indicating that they are determined by cell type identity.

We found that FACS-sorted Ch×10^+^ hindbrain neurons developed spontaneous rhythmic bursting activity *in vitro*. While there are no *in vitro* studies on unsorted hindbrain Ch×10^+^ neurons, this behavior is consistent with the observation that a closely related population of spinal V2a neurons develops spontaneous rhythmic activity following FACS isolation and reaggregation into three-dimensional cultures (Sternfeld 2017). Thus, FACS is a viable option for the isolation and subsequent long-term culture of molecularly defined neuronal subtypes.

### Cell type specific patterns of activity in cultures of sorted neurons

Several prior studies have arranged neurons on MEAs in very specific patterns (Maher 1999, Wheeler 2010), but not defined subtypes. The random patterning of molecularly defined cells on our arrays allowed us to explore whether there is a consistent influence of cell type on network behavior, regardless of network architecture.

Our observation that HB9^+^ motor neurons fail to develop spontaneous activity in the absence of glia is consistent with other studies that have demonstrated the essential support that astrocytes provide for cultured neurons (Wang 1994, Boehler 2007), including motor neurons (Ullian 2004). When we cultured sorted HB9^+^ neurons with astrocytes they developed unsynchronized spike trains. FACS appears to be critical for this behavior, as previous studies of stem cell-derived HB9^+^ motor neurons cultured without FACS isolation reported coordinated network bursts (Jenkinson 2017). In unsorted cultures, a cell type other than HB9^+^ neurons must have contributed to the generation of this activity pattern. We note that the spinal motor neuron differentiation protocol generates a small but prominent subpopulation of spinal V2a neurons, a cell type that is closely related to our rhythmogenic hindbrain Ch×10^+^ neurons (Brown 2014).

We found that Ch×10^+^ hindbrain neurons isolated by FACS and cultured on MEAs developed robust and highly coordinated network bursts. Calcium imaging (Figure 3) indicates that virtually all Ch×10^+^ neurons participate in these bursts, with no discernible time delay. Thus, simultaneous spiking is an intrinsic feature of Ch×10^+^ neurons in culture and does not appear to require the presence of other cell types.

### Ch×10-like pattern of activity is dominant in coculture

Recordings from the coculture indicate that Ch×10^+^ neurons impose their rhythmic bursting phenotype on adjacent HB9^+^ neurons (Figure 4e). In this way, we were able to induce HB9^+^ neurons to participate in network bursts by exposing them to another rhythmogenic cell type. Purified cultured HB9^+^ neurons did not coordinate their spontaneous calcium spikes despite being adjacent to each other. Thus, HB9^+^ neurons appear to require patterned input from another cell type. In contrast, Ch×10^+^ hindbrain neurons are able to generate their own patterns of activity without the need for exogenous cell types besides astrocytes. In summary, our results indicate that when cultured with astrocytes, electrically excitable cell types develop different spontaneous patterns of activity that are driven by the intrinsic properties of that cell type.

We were unable to detect any spontaneous activity in cultures without astrocytes, consistent with previous results with neuronal culture on MEAs (Ullian 2004, Boehler 2007). It is likely that one way that astrocytes mediate such an effect is by removing excess glutamate to prevent excitotoxicity (Rothstein 1996, Swanson 1997). Consistent with previous reports (Scemes 2006), the astrocytes in our culture were active, as indicated by slow waves of calcium activity which we were able to observe in calcium imaging (**Video 3-1**), but which did not produce electrical excitation on MEAs.

Our observation that Ch×10^+^ neurons are able to impose temporally patterned activity on HB9^+^ neurons is consistent with their *in vivo* function of driving rhythmic behaviors such as hindlimb locomotion and respiration. Prior studies suggest that activation of these neurons is associated with bouts of locomotion, and may drive locomotor stop signals (Bretzner, 2013, Bouvier 2015). Additionally, Ch×10^+^ neurons project to the pre-Bötzinger complex, and their ablation disrupts respiratory rhythms in newborn mice, with normal respiratory rhythms gradually reasserting themselves as the mice grow older (Crone 2012, Crone 2009).

### Emergent properties of neuronal cultures as revealed by synaptic inhibition

Our results from applying a panel of synaptic blockers targeting AMPA, NMDA, and GABA_A_ receptors to spontaneously active HB9^+^ and Ch×10^+^ neuron cultures (Figures 5 and 6) show that the AMPA_R_ blocker CNQX effectively blocked all bursts in Ch×10^+^ cultures and significantly decreased the activity in HB9^+^ neuron cultures. This is consistent with the observation that spinal motor neurons cultured *in vitro* form glutamatergic synapses that are entirely blocked by CNQX (Ullian 2004). CNQX application similarly eradicates spontaneous network bursting in cultures of spinal Ch×10^+^ neurons that are otherwise insensitive to glycine and GABA antagonists (Sternfeld 2017). Our finding that bursts of hindbrain Ch×10^+^ neurons could be effectively eradicated by blocking glutamatergic transmission suggests that the robust rhythmicity of these neurons is an emergent property of the network, as opposed to pacemaker activity generated by individual cells. This contrasts with true pacemaker neurons, such as those of the pre-Bötzinger complex, where bursts are intrinsic to individual cells, and therefore insensitive to the same cocktail of synaptic blockers (Chevalier 2016). Thus, we observe that AMPA receptor activation can drive very different outcomes that depend on cell type.

When we applied CNQX to the coculture, some neurons switched from rhythmic bursting to a transient period of tonic spiking before becoming quiescent. This emergent property may be driven by HB9^+^ neurons that revert to their native spiking phenotype in the absence of the patterning influence of network bursts. This is consistent with our calcium imaging data in which we identified HB9^+^ neurons in coculture that continued to have calcium spikes even in the presence of a dose of CNQX that effectively disrupted network bursts (Figure 5).

### Implications of our results for modeling reticulospinal circuits

The results of our study could be applied to modeling of reticulospinal circuits, different aspects of which are currently being examined by multiple groups (Sternfeld 2017, Oueghlani 2018, Pivetta 2014). In the rodent reticulospinal circuit, hindbrain Ch×10^+^ neurons primarily contact premotor networks within the spinal cord, as opposed to synapsing directly onto motor neurons (Bouvier 2015). By contrast, in the zebrafish hindbrain Ch×10^+^ neurons directly contact spinal motor neurons and generate swimming when selectively stimulated (Kimura 2013). Likewise, in *Xenopus* tadpoles the Ch×10^+^ dorsoventral hindbrain provides patterned excitatory input directly to motor neurons, driving sensory-evoked swimming before other motor control systems have developed (Soffe 2009, Li 2019). Thus, it can be argued that the circuit created by our *in vitro* cocultures replicates the basic circuitry found in fish and amphibians. It would be interesting to determine whether the emergent properties of Ch×10^+^ neurons from these species differ from the mouse, and how incorporating additional reticulospinal cell types would alter patterns of activity.

Ultimately, the most generalizable aspect of the findings we report here is the observation that the aggregate activity of neuronal networks is influenced by the specific molecular identity of their constituent neurons, beyond specific pacemaker cells or broad categories of excitatory-inhibitory cells. Our results indicate that many electrical properties of neurons are intrinsic to their specific subtype, which is an important consideration for modeling the effects of mutations and disease on network function. The cell type compositions of circuit models can have profound effects on patterns of activity and need to be considered and interpreted carefully.

## Acknowledgements

AVB, IT, LMK, and DWP designed research; AVB and HK performed experiments; AVB and IT wrote the paper.

We thank Selam Tadesse, Songyan Han, and Stanka Semova for technical assistance in flow cytometry experiments; The Rockefeller University Bioimaging Resource Center for microscopes; Carlos Rico for technical assistance with microscopy; Roger Vaughan for assistance in analyzing data; Kamal Sharma for providing Ch×10::CFP mice; Hynek Wichterle for providing HB9::GFP stem cells; and George Reeke, Bruce McEwen, and Mary Beth Hatten for their comments on the manuscript.

This work was supported by the New York Neuroscience Foundation, the Empire State Stem Cell fund through NYSDOH contract #C023046 and a Rockefeller Graduate Fellowship (AVB). IT was supported by NIH Grant F32 HD081835 and the Kavli Foundation. Opinions expressed here are solely those of the authors and do not necessarily reflect those of the Empire State Stem Cell Fund, the NYSDOH, the NIH, the Kavli Foundation, or the State of NY. All experiments on animals were approved by the Rockefeller University Animal Care and Use Committee. We declare no conflicts of interest.

## Supplemental data captions

**Video 3-1**: Calcium imaging of glia. Time course of Rhodamine3 fluorescence of astrocyte culture with spontaneous calcium activity. Playback of time 1-130s from raw recording (total time of raw recording is 130s). Minimum value of each pixel from the entire recording was subtracted from that corresponding pixel in each frame to minimize noise.

**Video 3-2**: Calcium imaging of spiking HB9^+^ neurons. Time course of Rhodamine3 fluorescence that corresponds to quantification in figure 3E. Playback of time 1-12s from raw recording (total time of raw recording is 80s). ROIs used to quantify HB9^+^ neuron calcium activity are outlined in yellow. Minimum value of each pixel from the entire recording was subtracted from that corresponding pixel in each frame to minimize noise.

**Video 3-3**: Calcium imaging of HB9^+^ neurons with coordinated spiking. Time course of Rhodamine3 fluorescence that corresponds to quantification in figure 3F. Playback of time 1-15s from raw recording (total time of raw recording is 80s). ROIs used to quantify GFP^+^ neuron calcium activity are outlined in yellow. Minimum value of each pixel from the entire recording was subtracted from that corresponding pixel in each frame to minimize noise.

**Video 3-4**: Calcium imaging of Ch×10^+^ neuron network bursts. Time course of Rhodamine3 fluorescence that corresponds to quantification in figure 3K. Playback of time 1-130s from raw recording. ROIs used to quantify Ch×10^+^ neuron calcium activity are outlined in yellow.

**Video 4-1**: Calcium imaging of HB9^+^/Ch×10^+^ neuron coculture. Time course of Rhodamine3 fluorescence that corresponds to quantification in figure 4E. Playback of time 40-80s from raw recording (total time of raw recording is 80s). ROI used to quantify HB9^+^ neuron’s calcium activity is outlined in green, ROI used to quantify Ch×10^+^ neuron’s calcium activity is outlined in cyan. Minimum value of each pixel from the entire recording was subtracted from that corresponding pixel in each frame to minimize noise.

**Video 4-2**: Calcium imaging of HB9^+^ neurons in HB9^+^/Ch×10^+^ neuron coculture. Time course of Rhodamine3 fluorescence that corresponds to quantification in figure 4F. Playback of time 1-80s from raw recording. ROIs used to quantify HB9^+^ neuron calcium activity are outlined in green. Minimum value of each pixel from the entire recording was subtracted from that corresponding pixel in each frame to minimize noise.

**Video 5-1**: Calcium imaging of bursting HB9^+^/Ch×10^+^ neuron coculture prior to AMPA_R_ blocker application. Time course of Rhodamine3 fluorescence that corresponds to quantification in figure 5H. Playback of time 35-75s rom raw recording (total time of raw recording is 80s). ROIs used to quantify HB9^+^ neuron calcium activity are outlined in green, ROI used to quantify Ch×10^+^ neuron’s calcium activity is outlined in cyan. Minimum value of each pixel from the entire recording was subtracted from that corresponding pixel in each frame to minimize noise.

**Video 5-2**: Calcium imaging of HB9^+^ neurons in co-culture after application of 40μM AMPA_R_ blocker. Time course of Rhodamine3 fluorescence that corresponds to quantification in figure 5I. Playback of time 8-38s from raw recording (total time of raw recording is 80s). ROIs used to quantify HB9^+^ neuron calcium activity are outlined in green, ROI used to quantify Ch×10^+^ neuron’s calcium activity is outlined in cyan. Minimum value of each pixel from the entire recording was subtracted from that corresponding pixel in each frame to minimize noise.

**Video 5-3**: Calcium imaging of bursting HB9^+^/Ch×10^+^ neuron coculture after washout of AMPA_R_ blocker. Time course of Rhodamine3 fluorescence that corresponds to quantification in figure 5J. Playback of time 10-40s from raw recording (total time of raw recording is 80s). ROIs used to quantify HB9^+^ neuron calcium activity are outlined in green, ROI used to quantify Ch×10^+^ neuron’s calcium activity is outlined in cyan. Minimum value of each pixel from the entire recording was subtracted from that corresponding pixel in each frame to minimize noise.

